# Synergism between Chromatin Dynamics and Gene Transcription Enhances Robustness and Stability of Epigenetic Cell Memory

**DOI:** 10.1101/2021.05.17.444405

**Authors:** Zihao Wang, Songhao Luo, Meiling Chen, Tianshou Zhou, Jiajun Zhang

**Affiliations:** Key Laboratory of Computational Mathematics, Guangdong Province; School of Mathematics, Sun Yat-Sen University, Guangzhou, 510275, P. R. China

## Abstract

Apart from carrying hereditary information inherited from their ancestors and being able to pass on the information to their descendants, cells can also inherit and transmit information that is not stored as changes in their genome sequence. Such epigenetic cell memory, which is particularly important in multicellular organisms, involves multiple biochemical modules mainly including chromatin organization, epigenetic modification and gene transcription. The synergetic mechanism among these three modules remains poorly understood and how they collaboratively affect the robustness and stability of epigenetic memory is unclear either. Here we developed a multiscale model to address these questions. We found that the chromatin organization driven by long-range epigenetic modifications can significantly enhance epigenetic cell memory and its stability in contrast to that driven by local interaction and that chromatin topology and gene activity can promptly and simultaneously respond to changes in nucleosome modifications while maintaining the robustness and stability of epigenetic cell memory over several cell cycles. We concluded that the synergism between chromatin dynamics and gene transcription facilitates the faithful inheritance of epigenetic cell memory across generations.

## INTRODUCTION

While cells carry information inherited from their ancestors and are able to pass on hereditary information to their descendants, they can also inherit and transmit information that is not stored as changes in their genome sequence. Such epigenetic cell memory, which is especially important in multicellular organisms, involves multiple biochemical modules, which can be roughly divided into three classes: chromatin organization [1], epigenetic modification [2] and gene transcription [3]. Each of these three modules can impact the other two. For example, covalent modifications at histone amino N-terminal tails can impact high-order chromatin conformation by facilitating the contact between histones and DNA or inter-nucleosomal interactions [4, 5]; In turn, spatial folding of chromatin is essential for enhancers to contact with distal specific promoters [6] and is also necessary for modified histones to spread their modifications to distant specific locus [7]. Chromatin dynamics (including chromatin organization and epigenetic modifications) can affect gene transcription and vice versa. For example, histone modifications occurring in the upstream area of a gene [8] can affect transcription factor (TF) access to regulatory sites [9] and further transcriptional initiation [10], thus impacting gene activity [11]. In turn, when bound to cognate regulatory sequences in gene regulatory elements, TFs either promote or suppress the recruitment of enzymes required for histone modifications [12] or chromatin remodeling [13]. In a word, relationships between the three modules are complex. Revealing these relationships is essential for understanding the robustness and stability of epigenetic cell memory.

Besides complex relationships, the three modules also behave on different timescales. Indeed, chromatin dynamic motion takes place on a timescale of seconds, whereas both epigenetic modifications and gene transcription occur on a timescale of minutes [14–21]. Thus, several important yet unsolved questions arise: whether or not there exists a synergetic interaction among the three modules, and how they dynamically collaborate to establish stable epigenetic memory patterns on multiple timescales, and what mechanisms govern the faithful inheritance of epigenetic memory over cell cycles.

Epigenetic modifications are essentially based on a “reader-writer-eraser” mechanism: functional enzymes “read” modifications and recruited enzymes then “write” spatially proximate histones [22,23]. Some studies showed that positive feedback loops [24] originated from the “reader-writer-eraser” mechanism, and long-range interactions [25,26] based on chromatin loops are essential for maintaining stable epigenetic cell memory [23,27,28]. Because of distinct chromatin conformations, there is a significant difference in the process that TFs search for a specific target position on DNA to regulate transcription initiation [29]. This fact, together with the evidence that histone modifications regulate 3D genome organization [4], suggests that chromatin modifications can predict gene expression [30]. Since the mechanisms underlying chromatin dynamics are possibly the ones for some part of the whole genetic and epigenetic regulation process, the synergic mechanism among the above three modules remain elusive.

Recently, mathematical models of chromatin dynamics, which are based on polymer physics but focus on the form of topological associated domains (TADs), were developed to explain how epigenetic cell memory is established and maintained [27,28]. However, TADs can partially reflect the relationship between chromatin conformation and epigenetic modification, and do not consider the dynamics of cellular life activities that affect epigenetic regulation. Analysis of other models involving gene expression and DNA replication important for cellular development and proliferation indicated that transcription and cell division antagonizing and perturbing chromatin silencing play an important role in stabilizing epigenetic cell memory [31]. In spite of their own advantages, the existing models of genetic and epigenetic regulations reveal neither the mechanism of how the above three modules collaborate nor that of how epigenetic cell memory is robustly and stably maintained due to this synergism.

In this paper, we propose a multiscale model to investigate the synergetic mechanism between a wide range of regulatory elements. Specifically, this model considers three correlated modules: one for 3D chromatin organization including various possible chromatin conformations, another for epigenetic modification including stochastic transitions between epigenetic states, and another for gene transcription including modification-mediated gene expression and transcription-regulated silencing antagonism. The first module, which is described by a generalized Rouse model, behaves on a fast timescale whereas the latter two, which are described by several biochemical reactions, behave on a slow timescale. Stochastic simulations of the multiscale model indicate that the epigenetic cell memory can be robustly and stably inherited through cell divisions. And the results reveal that the synergism among chromatin organization, epigenetic modification and gene transcription is essential for maintaining the faithful inheritance of epigenetic cell memory over generations.

## MATERIALS AND METHODS

### Modeling framework

Here we propose a strategy (in fact, a multiscale model) to model three coupled processes involved in gene expression: the formation of 3D chromatin conformations, epigenetic modifications, and gene transcription (Fig. 1). In the first process, chromatin is modeled as a chain consists of finite monomers (or beads) with each representing a nucleosome with a 3D position vector (Fig. 1a). This process also includes local and long-range interactions between nucleosomes. In the second process, epigenetic modifications including methylation and acetylation are classified as two different types: noise and recruitment modifications (Fig. 1b). The introduction of the third process is mainly because modifications affect transcription whereas transcription regulates silencing antagonism (Fig. 1c). Each of the three processes occurs on a different timescale. The elementary motion of chromatin is on a timescale from 10^−4^ to 10^−2^ (sec) [15]. Nucleosome modification dynamics based on the ubiquitous “reader-writer-eraser” mechanism is on a timescale of minutes [17,21]. And transcription occurring in discontinuous episodic bursts is on the timescale of about a minute (depending on regulation by enhancers) [18–20].

**Fig 1.**
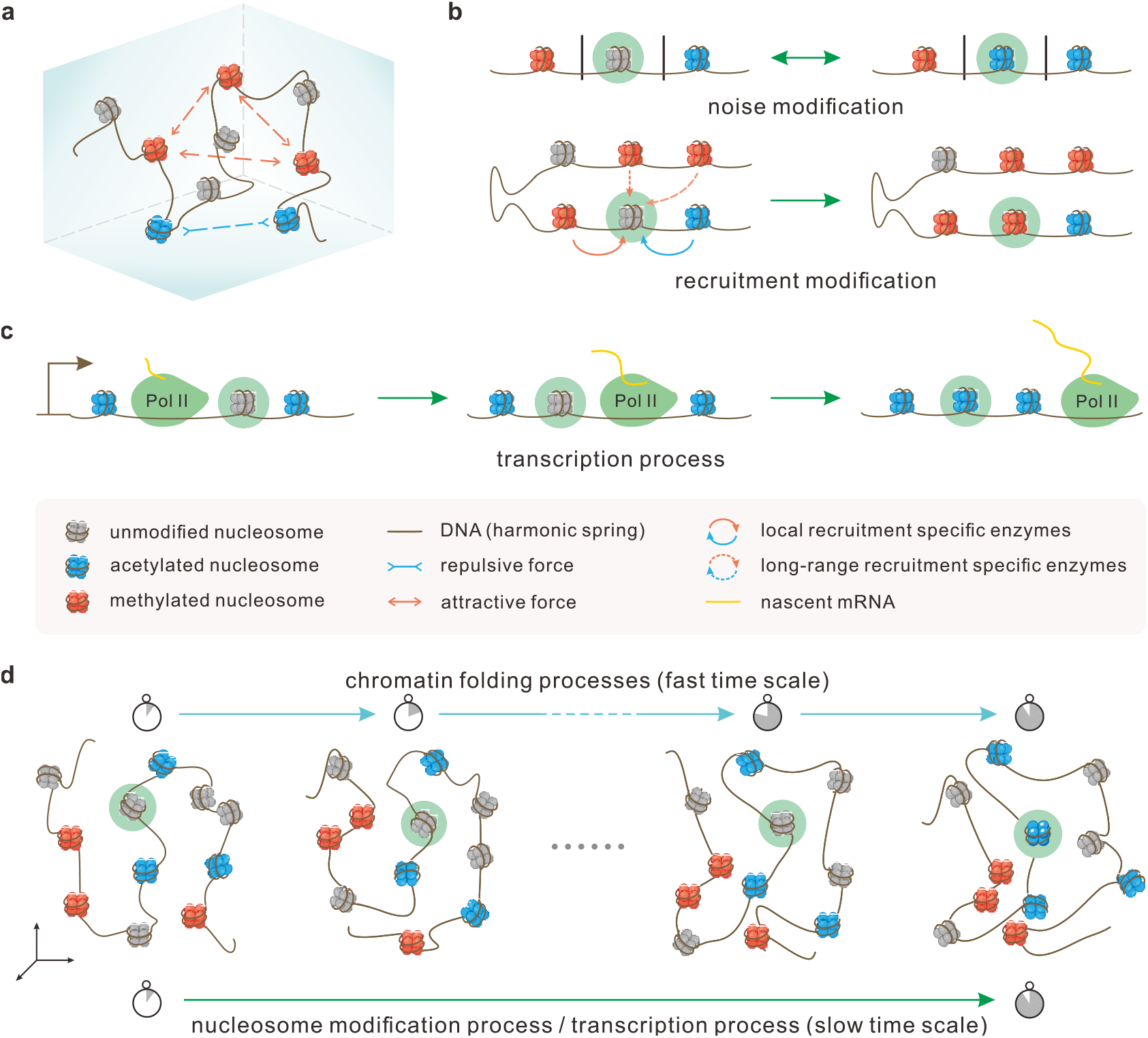
Schematic representation of a multiscale model that considers the relationship among three modules: chromatin organization, epigenetic modification, and gene transcription. **(a)** Schematic of 3D chromatin conformation. **(b)** Schematic of nucleosome modification regulation. Top panel: noise can induce a modification reaction. The modification state of a nucleosome is independent of its adjacent nucleosome states. Bottom panel: spatially adjacent nucleosomes have the ability to induce a modification reaction by recruiting specific modification enzymes. The farther away the nucleosome (marked by cyan shadow) is, the weaker is the effect. **(c)** Schematic of transcription process. Transcriptional state characterized by the presence of Pol II can drive nucleosome acetylation and demethylation. **(d)** Schematic representation of timescale differences among chromatin conformations, nucleosome modification and gene transcription in 4D space.

The above modeling strategy provides a possible framework for building 3D models and tracking cellular processes including transcription and cell mitosis over time (note: biological processes that rely on time-dependent dynamics is a 4D nucleome project [32,33]). Our multiscale model toward the 4D reality is a comprehensive investigation including the interpretation of mechanisms for the establishing and maintaining of stable epigenetic cell memory and the relationship between chromosome conformation, epigenetic modification and gene transcription. Although our model cannot accurately describe the reality of nucleolus epigenetic modifications in living organisms, it still captures the essential events occurring in gene-expression processes, including chromatin organization, epigenetic modifications, and gene transcription.

### Mathematical formulation

Chromatin is modeled as a polymer that is discretized into a collection of successive monomers connected by harmonic springs. Assume that the chain consists of *N* monomers. Each bead on the chain represents a nucleosome with the 3D position denoted by *P_i_* = (*x_i_*, *y_i_*, *z_i_*), where *i* = 1,…, *N*. We employ three kinds of multiple covalent modifications-acetylated (A, blue), unmodified (U, grey), and methylated (M, red) – to represent three possible epigenetic states of each nucleosome (Fig. 1), each denoted by *S_i_*∈{*A*,*U*, *M*}, where *i* = 1,…, *N*. Thus, (*P_i_*, *S_i_*) contains the position and modification information of the *i*th nucleosome. Since the multiscale model can be considered as the coupling of two different timescales - Brownian polymer dynamics of a fast variable and epigenetic modification and transcription of a slow variable, we adopt two distinctive yet correlative approaches to deal with the cases of the two timescales. Details are described below.

#### Fast time scale

The chromatin motion dynamics occur on a fast time scale. In our multiscale model, we use a generalized Rouse model with additional interacting beads (Fig. 1a and Fig. 2a) to describe the polymer structure. The conformational motion dynamics of the monomer *P_i_*(*i* = 1,…, *N*) is represented by the Langevin equation or the stochastic differential equation (SDE) of the form [34]

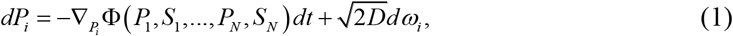

where Ф is the total potential of a given polymer conformation, *D* is the diffusion constant and *ω*_i_ is independent Gaussian noise with mean 0 and variance 1 in the 3D space. Chromatin structure dynamics evolve by the total potential Ф that will be specified afterward.

**Fig 2.**
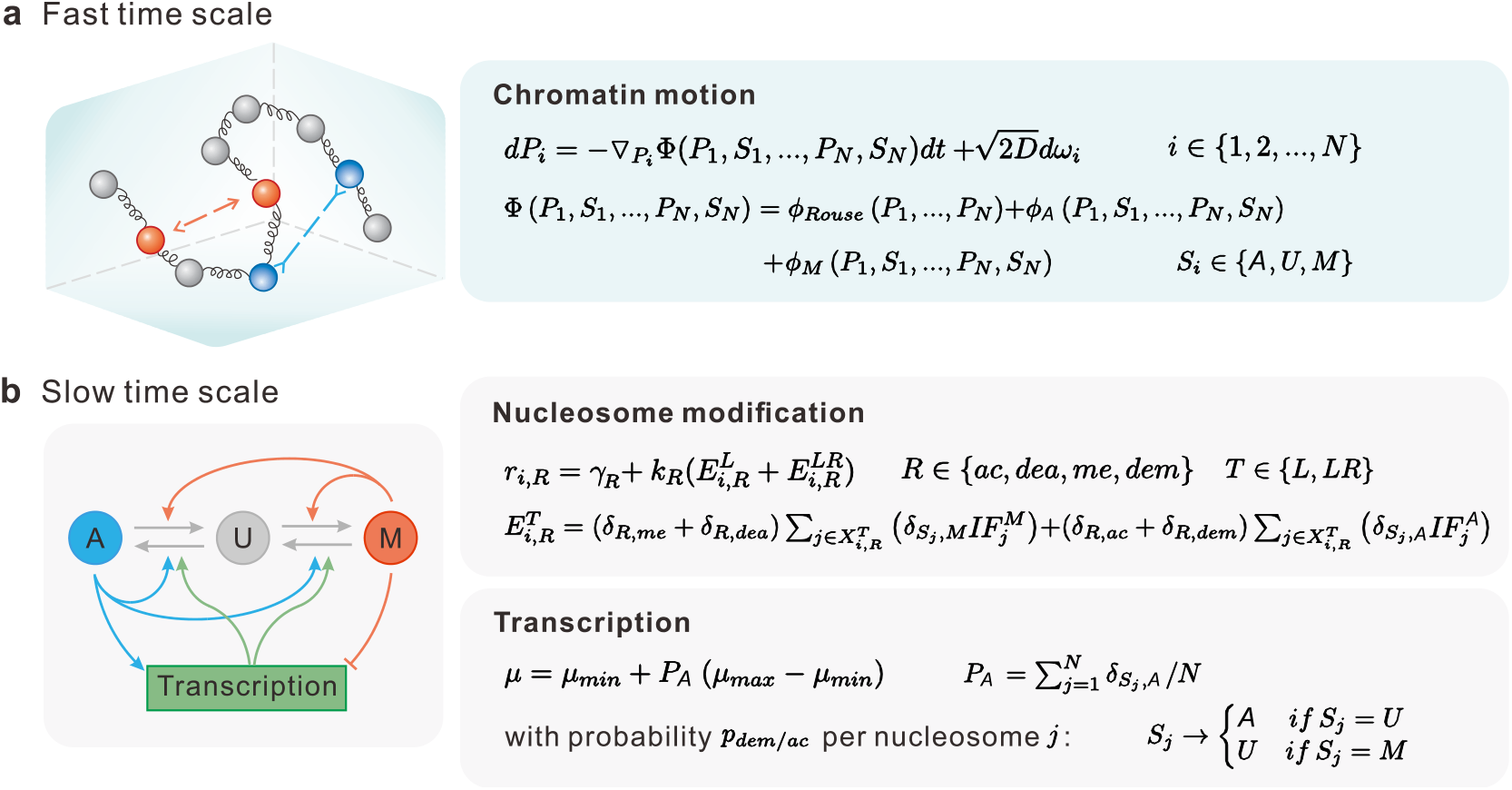
Mathematical representation of the multiscale model. **(a)** Mathematical model on a fast time scale. Left panel is a diagrammatic representation of the generalized Rouse model. Right panel is a mathematical equation of chromatin motion. **(b)** Mathematical model on a slow time scale. Left panel is a diagrammatic representation of feedbacks, where symbols A, U, M are referred, respectively, to acetylated, unmodified, methylated nucleosome state; grey arrows represent state conversions; colored arrows represent feedback interactions. Right panel is a mathematical formula for nucleosome modification and transcription. All symbols are described in the main text and Supplementary Table 1.

#### Slow time scale

Because of the fact that the chromatin conformation evolves much faster than nucleosome modification or gene expression (Fig. 1d), we can use a biochemical reaction system to describe slow variables in our model.

Each nucleosome in chromatin can be interconverted between epigenetic marks A, U and M. In general, a nucleosome with A (M) mark can be converted into M (A) state after the first mark has been removed to U (Fig. 2b) [23]. Each unmodified/modified process is considered a biochemical reaction. Thus, every nucleosome has four possible reaction channels - acetylation (ac), methylation (me), deacetylation (dea) and demethylation (dem). The corresponding biochemical reactions for the *i*th nucleosome (*i* = 1,…, *N*) read

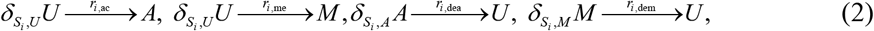

where *r_i,R_* is the rate of reaction *R* ∈{ac,me,dea,dem} for the *i*th nucleosome and will be discussed below, *δ*_*i,j*_ (Kronecker delta symbol) is equal to 1 if *i* = *j* and 0 otherwise. There are in total 4*N* reactions in our biochemical reaction system consisted of *N* nucleosomes.

Next, we consider transcription. An important point is that the multiscale model considers the relationship between transcription and chromatin epigenetic dynamics rather than the pathway how transcription occurs and how translational proteins act on chromatin. For simplicity, here we use a reaction to model transcription without considering the details of transcriptional burst on a slow timescale. This reaction reads

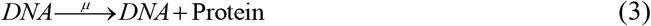

where *μ* is the mean transcription rate and will be discussed below.

The multiscale model framework defined by Eqs (1–3) is a coupled stochastic hybrid system. Previous studies have shown that an epigenetic regulation process is a complex and interrelated way [7,35], but the mechanism behind this process remains poorly understood. In contrast, our multiscale model proposes a feasible mathematical mechanism that, to some extent, can explain experimental phenomena and further draw qualitative conclusions as described in the abstract. Below we describe and elaborate on three different modules involved in our multiscale model.

### Three modules of the multiscale model

#### *Module 1*: 3D chromatin structure

It is the motion of nucleosomes that makes chromatin have distinctive 3D organization at the population level. Clearly, in our multiscale model described by Eq (1), the chromatin motion is mainly affected by the total potential Ф of a given polymer structure. In vivo, histone enriched in tri-methylations are linked to a higher condensed form of chromatin [36] and nucleosome acetylation state is associated with a less condensed organization [2]. Additionally, cell chemistry has shown that methylations do not change the charge of residues; yet, they alter the overall size of the modified amino acid residues. In contrast, acetylated nucleosomes have the ability to neutralize the positive charge of amino acid, thus inducing a less condensed conformation. These facts imply that the presence of different nucleosome modifications would have an important effect on the structure of chromatin. Presumably, besides the effective potentials between consecutive monomers, we add interaction forces mediated by the epigenetic marks in the generalized Rouse model: methylated marks attract to each other; acetylated marks are mutually repulsive; there is no interaction between unmodified monomers, but they can participate in epigenetic modifications (Fig. 1a).

Thus, for a given conformation and epigenetics of a polymer, the total potential may be represented by

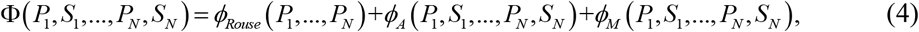

Where 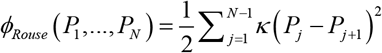 is an effective potential between consecutive monomers, κ is the stiffness of the spring, 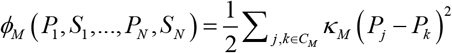 and 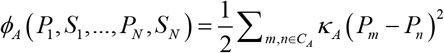 are energy potentials of methylated and acetylated monomers respectively, *κ _M_*, *κ _A_* are the attractive and repulsive interaction coefficients, *C_M_* and *C_A_* are the ensembles of indices for nucleosome methylation and acetylation. More details on the generalized Rouse model are given in Supplementary Note 2.

#### *Module 2*: Transitions between epigenetic modification states

Each nucleosome can dynamically transition between epigenetic marks A, U and M, according to Eq (2). In the following, we define the rates of biochemical reactions for transitions from two perspectives (Fig. 1b):

##### I. Noisy modification

Nucleosomes can be interconverted by noisy modification (corresponding to non-feedback processes), which is primarily due to the leaky enzymatic activity or the effects outside the region boundaries. More precisely, the nucleosome modification status is independent of the adjacent nucleosome states. We assume that the noisy modification rate of each nucleosome takes a constant or a certain proportion of recruitment modification. Specifically, we set noise rates *γ_R_*, *R* ∈{ac,me,dea,dem} at 5% [31] of the rate corresponding to recruitment modification *k_R_*.

##### II. Recruitment modification

Nucleosomes can also be interconverted by recruitment modification (corresponding to feedback processes), which is related to the propagation of the epigenetic mark by recruitment of the enzymes corresponding to other locus. This process [37,38] forms positive feedback loops in the reaction scheme: nucleosomes with A or M modification recruit protein complexes to promote spreading of the state or erasing of the antagonistic mark. Here, we assume two types of feedbacks: (**a**) methylation (acetylation) state can promote the process from un-modification to methylation (acetylation); (**b**) methylation (acetylation) state can promote the process from acetylation (methylation) to un-modification (Fig. 2b). Yet, the mechanism and relationship between these two types of feedbacks are not clear. We hypothesize that the latter has a 10-fold reduced efficacy of the former [31], that is, *k*_dea_ = *k*_me_/10, *k*_dem_ = *k*_ac_/10.

For the *i*th nucleosome, the spatial adjacent modified nucleosomes can participate in its recruitment modification, but the efficacy of modification decreases with increasing nucleosome separation [23,39]. We call the magnitude of modification efficiency an impact factor. There are two types of impact factors: the set *X_i_* of methylated (or acetylated) nucleosomes around the *i*th nucleosome affects its acetylated process to its methylated process or vice versa. Note that each methylated (or acetylated) nucleosomes in *X_i_* has a corresponding impact factor 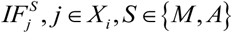, *j* ∊ *X_i_*, *S* ∊ {*M, A*} on the *i*th nucleosome (see discussions below). In fact, the value of impact factors is related to the structure of chromatin. Thus, we consider the spatial position of nucleosomes and decompose the recruitment modification in two distinctive contributions:

###### (i) Local interaction (L)

Modification of a nucleosome is constrained to spread through its two nearest neighbors on the polymer chain. As shown in Fig. 1b (solid line), the enzymes recruited by the left or right nucleosome can work on the middle nucleosome. Certainly, such a restriction might also arise through steric limitation, which exists merely when adjacent nucleosomes meet. For each nucleosome, the impact factors of the left and right nucleosomes are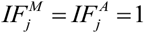

##### (ii) Long-range interaction (LR)

Chromatin motion including chromatin loops that bring distant loci into close spatial proximity [7] can form effective long-range interactions [24]. For a nucleosome, the nucleosomes in its adjacent spatial neighbors, not merely its nearest-neighbors along the chain, recruit specific enzymes and affect its change (Fig. 1b (dashed line)). In our model, we use the contact probability of two nucleosomes in space to approximately reflect the effectiveness of modification. Thus, we can assume that when the spatial distance between nucleosomes exceeds a certain value, the impact factor decreases, which is usually represented by a power law ∝ *d*^−3/2^[40], where *d* is the separation distance in the 3D space rather than the genomic distance. In addition, we know that higher methylation indicates increased chromatin compaction but higher acetylation expresses reduced chromatin compaction. Therefore, it is reasonable to set different spatial interaction gyration according to different modifications: acetylated (methylated) monomer has a larger (smaller) interaction threshold. The impact factor can thus be expressed as

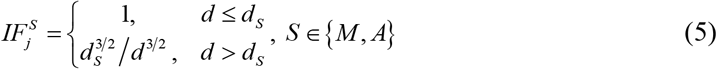

where *d_S_* is the threshold of spatial interaction distance of the nucleosome in *S* state, *d* is the spatial distance between two nucleosomes.

Putting those together, the rate for the reaction *R* ∈{ac,me,dea,dem} for the *i*th nucleosome (*i* = 1,…, *N*) is

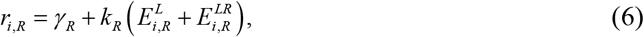

where

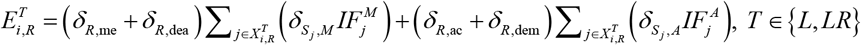

is the sum over local and long-range interacting nucleosomes, and 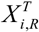 is the set of local and long-range interacting nucleosomes that recruit the corresponding enzymes to affect monomer *i* in reaction *R*, and *γ*_R_ is the noise modification rate and *k_R_* is the recruitment modification rate.

#### *Module 3*: Modification-mediated gene expression and transcription-regulated silencing antagonism

There is evidence to support that TFs regulate gene expression partially by nucleosome modifications [41,42]. However, the mechanistic basis of transcription dependence of modification levels remains an open challenge. From experimental observations and previous models, we know that A is an open conformation that the gene promoter is accessible to TFs and conducive for transcription [2] (e.g., acetylated H3K9, H4K16), and M is the repressed chromatin state that is assumed to be related to silencing [43,44] (e.g., methylated H3K9, H3K27) although not all methylations suppress gene expression (or transcription) [45]. The reason we make this assumption is that methylation and acetylation on the same nucleosome, such as H3K9 and H3K27 have distinct states, thus it is convenient to compare methylation with acetylation. Additionally, we hypothesize that the level of RNA production depends on the methylation or acetylation level. And the initiation rate of transcription *μ* is a simple linear function of the proportion of the number of acetylated nucleosomes, that is,

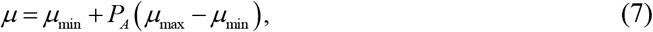

where 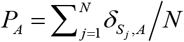 is the proportion of acetylated marks, *μ_min_* (*μ_max_*) is the minimum (maximum) transcription initiation rate. Note that the 3D chromatin shape also impacts the gene activity by limiting the accessibility of Pol II. The reason why we use epigenetic modification to measure transcriptional activity is that transcription is measured more accurately by modification on a slow timescale than by structure on a fast timescale. Structure and modification of chromatin are directly related as discussed above, so we assume that the interaction between structure and transcription is reflected in modification.

In addition, demethylation is associate with the fact that demethylase is located in the promoters and the coding regions of protein complexes for target genes [46,47]. Therefore, when transcription occurs, this promotes the transition of methylation state to an unmodified state [48]. Moreover, there is evidence to support that protein complexes involved in transcriptional activation lead to the identification of a large number of histone acetyltransferases [5], which can enhance the conversion of unmodified state to deacetylation state. Considering all the above facts, we model transcription as directly antagonizing epigenetic silencing [49,50] that causes removal of M state or add of A state (Figs. 1c and 2b). Therefore, we posit that each transcription is viewed as a discrete event that causes nucleosome demethylation and acetylation with probability *p_dem/ac_*.

### Simulation method and statistics

According to the above description, we use the above SDE (i.e., Eq. (1)) to simulate chromatin motion on a fast time scale and Gillespie stochastic algorithm [51] to simulate biochemical reactions (i.e., Eqs. (2) and (3)) on a slow time scale. The latter generates an exact pathway *a* and a time step *τ* in the light of a number of reaction channels and the corresponding propensity functions. At each iteration, *τ* corresponds to the typical time-scale of modeling (see Supplementary Note 2). We hypothesize that the chromatin structure and modification state of the current moment determines when the next reaction occurs and which reaction will occur according to the Gillespie algorithm, as well as when the system time reaches the moment of the next reaction occurring so that the specific enzyme promotes the selected reaction (Fig. 1d and Supplementary Fig. 2). In addition, we use 10^−2^ (sec) to represent the time step of chromatin folding.

When the cell reaches the end of the cell cycle with a timescale of 22 hours [52], DNA replication and cell division occur. We assume that de novo nucleosomes participate in the two copies of DNA, and both old and new nucleosomes are normally shared at random between two daughter chromosomes [53]. With this assumption, each nucleosome is replaced with a new unmodified nucleosome with a probability of 0.5. Numerical simulations are performed using a home-made program (written in MATLAB). The whole system is simulated according to the flowchart in Supplementary Fig. 3. Snapshots of the system are taken every 300 seconds. Using the generated data, we then carry out a quantitative analysis.

**Fig 3.**
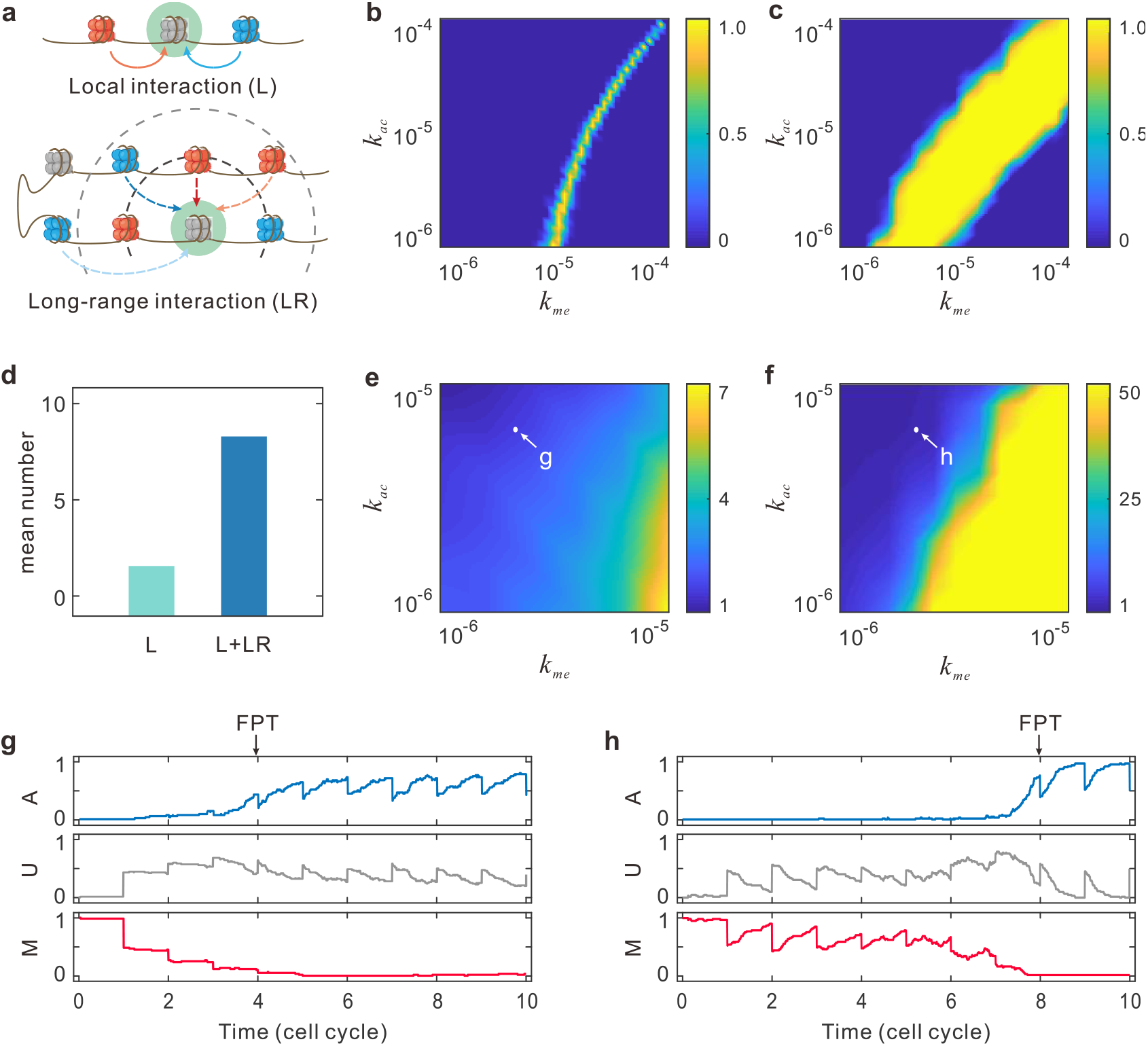
Effect of chromatin organization on epigenetic cell memory. **(a)** Schematic representation of two distinctive interactions. Top row: local interaction, where modification of a nucleosome spreads through its two nearest neighbors along the chain. Bottom row: long-range interaction, where modification of a nucleosome spreads through its adjacent spatial neighbors. The impact efficacy decays with increasing spatial distance (with a lighter dashed line). **(b)** Heatmap, showing that bistability measured by quantity *B* is taken as a function of methylation rate *k*_me_ and acetylation rate *k*_ac_ without long-range interaction. For each set of parameter values, 100 simulations are initialized in each of the uniform methylation or acetylation states, and 50 cell cycles are considered. Results are obtained by averaging over all simulations. **(c)** Heatmap similar to (b) but for long-range interaction. **(d)** The average number of modified nucleosomes around an unmodified nucleosome under the situation that long-range effects are present or absent. **(e)** Heatmap for the local spreading model of the MFPT *tFP*(*M*) for switching from the methylation state to acetylation state as a function of *k*_me_ and *k*_ac_, which is obtained by averaging over 100 simulations for each of 50 cell cycles. **(f)** Heatmap similar to (e) but for the long-range spreading model, where two points indicated by (g) and (h) are used in detailed analysis. **(g)** An example for stochastic simulation of the levels of modified and unmodified nucleosomes with initial uniform methylated state overtime for an acetylation-biased local spreading model, where parameter values are *k*_me_= 2×10^−6^ and *k*_ac_ = 6×10^−6^. **(h)** An example similar to (g) but for an acetylation-biased long-range spreading model.

The global epigenetic state is measured by calculating the epigenetic magnetization

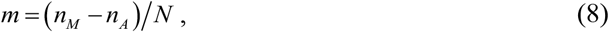

where *n_M_* (*n_A_*) is the number of methylated (acetylated) nucleosomes in the system. At a high magnetization, chromatin is filled with methylated nucleosomes so that the chromatin is dense. Pictorially, the radius of gyration takes the form

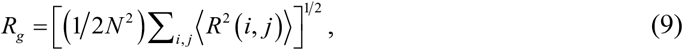

where *R*^2^(*i*, *j*) is the pair-wise squared distances. The radius defined in such a manner can characterize the looseness of polymer in 3D space: It has a higher value when the polymer is an open (acetylated) conformation, and a lower value when it becomes a compact (methylated) globule.

Because of considering gene-expression reactions without considering the details of transcription and translation, we count the number of times for the occurrence of transcription in the interval of snapshots as the feature of gene activity.

Putting all the above details together, we have a novel theoretical model in which epigenetic gene regulation and chromatin architecture are mechanistically integrated on different timescales. This 4D multiscale model actually gives a method of mapping the structure and dynamics of chromatin in space and time, thus gaining deeper mechanical insights into how epigenetics is maintained after several cell cycles and what mechanisms enable the three modules to work together dynamically. More details of the model and values of the parameters used in the simulation are given in Supplementary Information.

## RESULTS

To explore the power of the above multiscale model in painting 1D epigenetic information, 3D chromatin structure and gene transcription, we examine a system consisting of *N* = 60 nucleosomes that corresponds, typically, to a small domain (~12 kb of DNA). To simplify, the simulation region is isolated from neighboring the DNA by boundary insulator elements [54,55].

### Chromatin organization driven by long-range interaction can enhance epigenetic cell memory and its stability

Note that our model explicitly considers the chromatin spatial structure and the accurate dissection of the 3D contributions to nucleosome modifications (Eqs. (2) and (6)) (Fig. 3a). Therefore, the questions we first want to answer are what role chromatin organization plays in maintaining epigenetic cell memory in the sense of gene expression and stability of chromatin status, and how chromatin folding properties affect epigenetic processes. For this, we consider a controlled system without long-range epigenetic modifications. In other words, the dynamics of epigenetic modification do not depend on the folding of the chain, and the question thus reduces to a simpler 1D one. For this controlled system, Eq. (6) reduces to 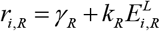.

To maintain epigenetic cell memory in the sense of gene expression, the modified chromatin must have the ability to maintain epigenetic patterns - high acetylation (low transcription) and high methylation (high expression) states - for several cell cycles. If a model is capable of sustaining both high-M state and high-A state under the same conditions, it is bistable. In order to better characterize this property, we perform a set of simulations over a range of parameter values to calculate the balanced bistability *B* = 4*P_M_P_A_* [56] (see Supplementary Note 3), where *P_M_* or *P_A_* is the probability that the system is in one of the epigenetic states. If *B* approaches to 1, the system is bistable.

By simulations, we find that the system exhibits weak bistability without long-range epigenetic modifications (Fig. 3b) but strong bistability with both local and long-range interaction (Fig. 3c). If either methylation rate *k_me_* or acetylation rate *k_ac_* is much larger than the other, the system cannot exhibit bistability. With increasing acetylation rate *k*_ac_, the minimum *k_me_* for bistability is observed to increase. This is because the replacement rate of the methylated nucleosome is not quickly enough to counteract demethylation, acetylation, transcription and DNA replication. Since transcription antagonizes silencing, this promotes the process from acetylation to methylation. Therefore, the bistability is observed almost in the region where the methylation rate is bigger than the acetylation rate. In short, if the process from acetylation to methylation and the process from methylation to acetylation can be balanced, the system can be bistable, indicating that the chromatin can store both active and repressive epigenetic memory, and inherit epigenetic states for several cell cycles.

We calculate the average number of the modified nucleosomes that can influence modification around an unmodified nucleosome (Fig. 3d). Then, we find that without long-range epigenetic modification, the average number is 1.5, and the number for 3D chromatin is as high as 8.5. This implies that the long-range interaction brings about seven modified nucleosomes caused by the chromatin motion as shown in Fig. 3a. Thus, we can draw the conclusion that the structure of chromatin has a great influence on the modification of nucleosomes by enhancing and reinforcing epigenetic cell memory.

Let us explain the effect of chromatin folding from another more intuitive perspective. For this, we simulated our model using different values of methylation and acetylation rates starting from the initial repressed state, and calculated the mean first passage time (MFPT) for switching from a M macro-state *m* = 1 to an A state (*m* < 0) (see Supplementary Note 3). Fig. 3e and Fig. 3f show the heatmaps of the MFPT in the presence and absence of long-range effect, respectively. We observe that a larger methylation rate leads to a higher MFPT. On the contrary, a larger acetylation rate results in a lower MFPT. Moreover, the epigenetic system is extremely unstable without the long-range crosstalk (Fig. 3e). The introduced-above effective long-range interaction can stabilize large-scale epigenetic states dramatically (on average, more than 4-fold compared to the case of the local spreading model) (Fig. 3f). By comparing Fig 3c and Fig. 3f, we find that in 50 cell cycles methylation state does not switch to acetylation state in the bistable region, showing the robustness of the epigenetic cell memory. Fig. 3g shows the result of a stochastic simulation without long-range interaction, whereas Fig. 3h shows the result of another stochastic simulation with long-range interaction. In the two cases, the parameter values are the same, but the repressed state is erratic and biased to the acetylation state for several cell cycles. We can see that the epigenetic cell memory is more unstable without long-range interaction, and the shorter time for transiting to an active state. This means that the 3D spreading of a mark leads to the spontaneous formation of a more stable epigenetic coherent phase, implying that 3D chromatin conformations are important for stabilizing epigenetic heritage.

### Chromatin structure and gene activity can promptly and simultaneously respond to changes in modification

Here we examine the effect of directly changing the modification rates on chromatin states. Our strategy is that we first simulate the model with *k*_me_ = 7×10^−6^ and *k*_ac_ = 7×10^−6^, starting from the active state after equilibration for ten cell cycles, and then change *k*_ac_ to observe the epigenetic kinetics (Fig. 4a and Fig. 4b). We observe that if the alternation of *k*_ac_ is small, e.g., *k* =1×10^−6^ (Fig. 4a), the epigenetic state does not change but produces controllable fluctuations, indicating the robustness of stable memory. If *k*_ac_ is altered to *k*_ac_ =1×10^−7^ at *t* = 0, Fig. 4b shows a simulation where the stable high A modification coverage becomes unstable and biases toward M modification.

**Fig 4.**
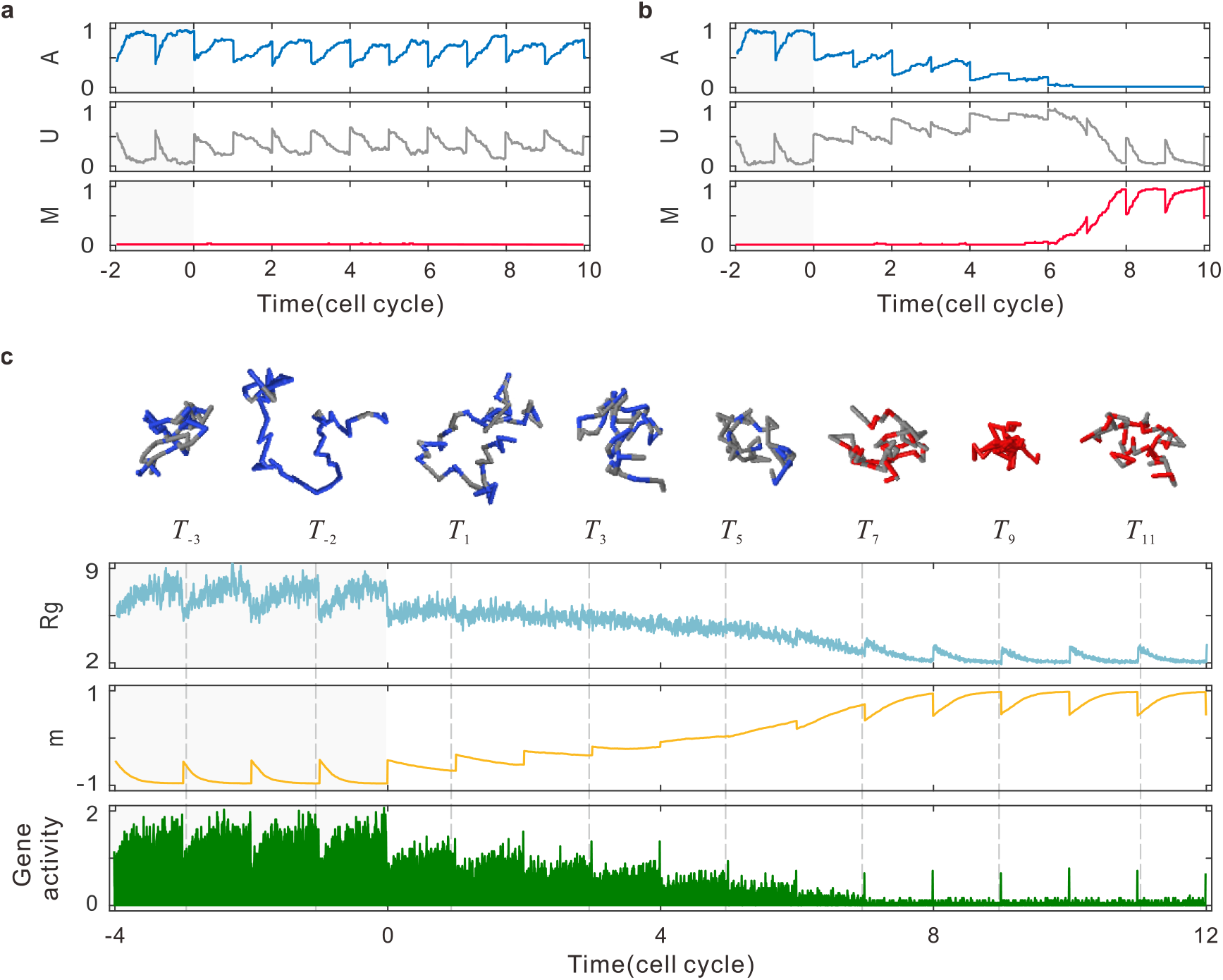
Responses of chromatin structure and gene activity to changes in nucleosome modification. **(a)** The time evolution of the levels of modified and unmodified nucleosomes, where after initialization of uniform methylated state for ten cell cycles (two of them are shown) with *k*_me_ = 7×10^−6^ and *k*_ac_ = 7×10^−6^, parameter *k*_ac_ is changed to *k*_ac_ =1×10^−6^ at time *t* = 0. In spite of this perturbation of *k*_ac_, the chromatin state are still maintained for several cell cycles. **(b)** Except for *k*_ac_ =1×10^−7^ at *t* = 0, the other parameter values are kept the same as (a). Following this perturbation, the chromatin state turns to the active state and persists for several cell cycles. **(c)** Top row: Typical snapshots of 3D structures by once simulation, where the polymer is taken as a function of time, *T_i_* represents a certain time in the *i*th cell cycle, *T*_−3_ and *T*_11_ represent the initial stages of cell cycles and other *T_i_*s represent the final stages. Bottom rows (2-4): The time evolution of the average values of radius gyration *R_g_*, epigenetic magnetization *m* and gene activity with multiple simulations. Conditions change but parameter values are the same as (b).

Clearly, we should consider the impact on structure and gene activity under the chromatin state transition rather than under the stable state. Additionally, since there are fluctuations and differentiations in gene-expression frequency and chromatin size for once simulation and the FPT of each simulation is different, we simulate several times and then select simulations with roughly the same FPT and calculate the mean value of the radius of gyration *R_g_*, magnetization *m* and gene activity, which can be represented as the characteristics of structure, epigenetic, and function of chromatin, respectively.

At the initial four cell cycles in Fig. 4c, the *R*_g_, *m* and gene activity is regular with the time evolution. In each cell cycle, the magnetization is about −0.5 because of dilution at DNA replication at the beginning, and is then reduced to −1 gradually with the spread of epigenetics. Meanwhile, the chromatin structure is gradually slackening to facilitate the binding of transcriptional enzymes. Thus, the expression of the gene is stably active and the level of acetylation determines the level of the expression according to Eq (7).

We can see that if the parameter *k*_ac_ is altered to *k*_ac_ =1×10^−7^ at *t* = 0, the *R_g_*, *m* and gene activity responds, immediately, to the alteration of the epigenetic rate. At the end of the 1^st^ cell cycle that has changed the rate, the *m* does not recover to −1, even though the number of acetylated nucleosome has a slight increase (Fig. 4b) due to the effect of positive feedback loops of acetylated states and the effect of transcription (Fig. 2b). Meanwhile, in one cell cycle, the *R_g_* and gene activity is also responded promptly (Fig. 4c).

The cell has the tendency to be methylated and turns to steady high M coverage at the 8^th^ cycle (Figs. 4b and 4c). The intermediate part can be viewed as a short window, in which the system can switch from high A to high M modifications. We can see that in this window, the amplification of M on the whole is simultaneously accompanied by the decrease of *R_g_* and gene activity in multiple cell cycles (Fig. 4c). At the beginning period of the window, the number of acetylated nucleosomes is decreasing with increasing unmodified in multiple cell cycles. In the latter period of the window, the number of acetylated labels decreases drastically, triggering a rapid rise of methylation. Thus, the *m* is progressively increasing and clusters of methylated modifications emerge at the end of short window. At the same time, we can find that *R_g_* gradually attenuates and the chromatin condenses fairly slowly, which persists for several cell cycles. Fig. 4c shows typical snapshots of 3D shapes, which entirely display the switching process from state A to M. On average, the gene activity is gradually decreasing due to the level of acetylation and spatial condensing of chromatin (Fig. 4c).

When the system turns to steady methylation, the gene is almost silent through the whole cell cycles due to little acetylation. Moreover, in a cell cycle, the methylated nucleosomes accumulate, resulting in an increasing global epigenetic modification and a decreasing radius of gyration.

In summary, we find an interesting phenomenon: the number of acetylated nucleosomes is decreasing with reducing transcription probability and shrinking the radius of gyration, and vice versa. This phenomenon would imply that a cell has the ability to alter its state in response to external changes and that chromatin structure and gene activity can simultaneously and immediately respond to the changes in modification rates.

### A synergetic self-organization strategy for genetic and epigenetic regulations

The above results indicate that three modules of our model – dynamic spatial motion of chromatin, stable epigenetic modification and genetic function of chromatin - can simultaneously make dynamic and timely adjustment, in face of internal and external noise from, e.g., alterations in modification rates. This suggests that the synergy among these three modules can regulate genetic and epigenetic processes.

In order to explain the possibility of such a synergy or the rationality of such a strategic mechanism, we simulate a wide range of parameters of *k*_me_ and *k*_ac_ over 100 simulations for each of 50 cell cycles, and record the nucleosome position and modification information at the end of each cell cycle and the mean gene activity in the last hour of each cell cycle. Then we calculate the Pearson correlation coefficients between the radius of gyration *R_g_*, magnetization *m* and gene activity. Note that this coefficient describes the covariation of two random variables, and takes a value between −1 and 1.

We observe a strong positive correlation between chromatin organization and expression level (Fig. 5a, *r* = 0.65) and a strong negative correlation (Fig. 5b, *r* = −0.73) between chromatin structure and nucleosome modification level. High coefficients represent that the information of chromatin organization is promptly transferred to the gene function and chromatin modification to adjust the epigenetic process. In the cases of long-range interaction and no long-range interaction, Fig. 3 has partially shown that the information on chromatin structure can affect the modification process. Two strong negative correlations in Fig. 5b and Fig. 5c (*r* = −0.88) suggest that a different nucleosome modification level can induce a distinct gene expression pattern and adjust the spatial folding of epigenomes simultaneously and promptly. Fig. 4 has shown a complete process from stable acetylation to stable methylation, which is caused by the alteration of modification rate and is accompanied by the changes of gene activity and organization. Fig. 5a and Fig. 5c also suggest that the transcriptional events can influence nucleosome modification and chromatin structure. In multicellular organisms, these correlations might be derived from a variety of enzymes in the cellular activity. In fact, an enzymatic reaction or a binding behavior trigger a cascade of molecular events that affect the function or action of the cell. Thus, we can conclude that in the combination modeling of fast and slow time scales, any two of the three modules are correlative and even strongly correlative as shown in Fig. 5. These correlations occur due to the effect of the long-range interaction, but they will become weaker if the long-range interaction is not considered (or if the only local interaction is considered) (referring to Supplementary Fig. 4). Therefore, we conclude that the long-range interaction rather than the local interaction is a key factor for the synergism among chromatin structure, epigenetic modification and gene activity to maintain the stable epigenetic cell memory.

**Fig 5.**
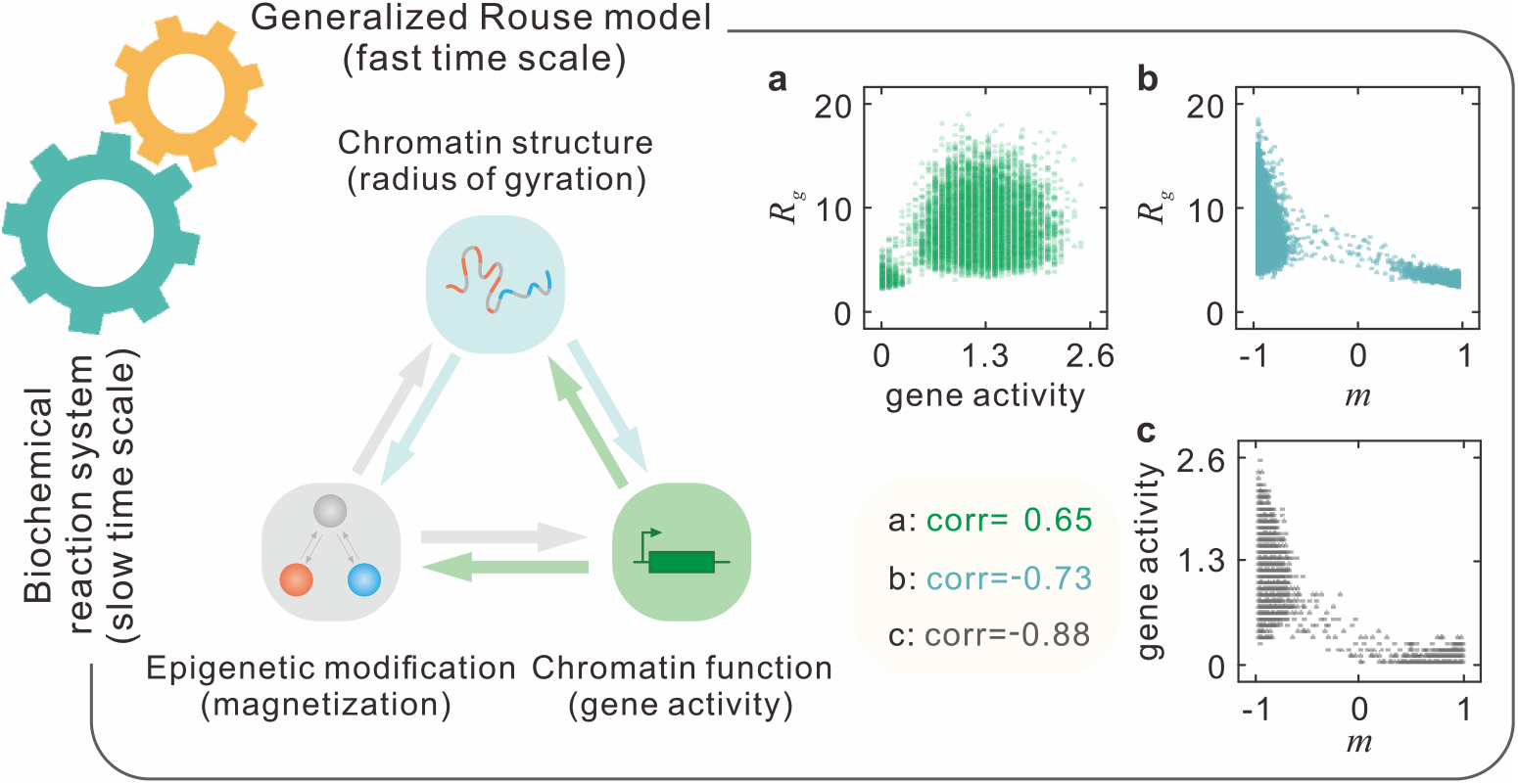
**A multiscale dynamical model for the synergism among chromatin structure, epigenetic modification and gene activity**, where “corr” represents the Pearson correlation coefficient between two random variables.

In addition, in our multiscale model, the associated methylation or acetylation labels favor chromatin self-attractive or self-repulsive interactions, and these, in turn, drive the formation of distinct structure through updating the energy potential of the system. These different conformations might influence the communications of different marks via long-range interaction and the diffusions of specific enzymes binding to enhance transcription. According to cell biology and biophysics, we know that the alternative modified nucleosomes suffering the positive feedback mechanism result in a regulated expression process through tuning transcription rate. In turn, transcription urges the transition from methylation to acetylation via discrete turnover events in order to sustain the positive feedback of gene activity, and further drives the folding of the polymer to some extent. Finally, the interaction between chromatin structure and gene transcription is reflected by modification, not only because they are all related to modification but also because transcription and structure are at different timescales. It should be pointed out that the described-above relationships among chromatin structure, epigenetic modification and chromatin function hold for multi-generations. Therefore, we can conclude that the synergism among the three modules shapes a stable genetic and epigenetic network (Fig. 5).

## DISCUSSION

In this paper, we proposed a multiscale stochastic model to investigate the robustness and stability of epigenetic cell memory. This model focuses, especially, on the cooperative interaction among chromatin spatial motion, stable epigenetic modifications and chromatin genetic function (in fact, gene activity). It provides a formalism of realistic biological processes in which enzyme modifications and transcription occur on a slower timescale than chromatin spatial folding. In spite of the difference in timescale, the mentioned-above modules can collaborate (Fig. 5) to drive and even control cell fate determinations through a stable genetic and epigenetic networks.

Previous studies showed that long-range epigenetic modifications can facilitate nucleosome-nucleosome communication and histone modification propagation [7], and control gene expression [57]. For example, an acetyltransferase is recruited to the enhancer, which triggers the increase of H3K27 acetylation at the promoter and subsequent transcription [58]. And in *Drosophila melanogaster*, temporal and spatial expression of Hox genes during development depends on Polycomb group proteins and on the long-range contacts between the Hox locus and distal specific enhancers [59]. In contrast, here we have shown that the long-range interaction can reinforce the stability and robustness of epigenetic cell memory over several cell cycles.

How chromatin state and gene activity respond to changes in epigenetic modification is a fully unsolved issue in the field of molecular biology. First, cells have the ability to sense and adapt to environmental changes. Second, small external noise is not sufficient to destroy epigenetic cell memory due to the coherent formation of epigenetic modification. This machinery may endow regulatory networks with enhanced robustness. However, when external noise is large such as climate cycle in spring or winter and artificially increased enzyme concentrations in experiments, the chromatin-based noise filtering machinery cannot completely eliminate the noise impact. Thus, jumping into an alternative landscape epigenetic state due to the noise effect will occur with a large probability (this corresponds to the plasticity of cells [60]), e.g., vernalization in *Arabidopsis* centres on the *FLC* gene [61]. Exposing to the prolonged cold of winter, the *FLC* gene, a repressor of flowering, fills with acetylated nucleosomes [62,63], and after vernalizaiton, the expression of *FLC* is stably repressed and the plants has the ability to flower with the modification biasing towards methylation. In our modeling framework, changes in modification rates or other parameters such as the transcriptional initiation rates model can be considered as exogenous stimuli. Thus, our model has plasticity and extendibility. Moreover, we have shown that chromatin state and gene activity respond, promptly and simultaneously, to changes in modification rates.

The synergism among chromatin organization, histone modification and gene transcription is critical for the maintenance of stable epigenetic cell memory. For example, when the β-globin locus is located in a highly acetylated environment, it will increase the sensitivity to DNase so that the chromatin structure can have universal accessibility [64]. The synergism is also important for chromatin states switching in face of complex external environments, e.g., the vernalization in *Arabidopsis* centres on the *FLC* gene discussed above. However, if the synergism is broken or if any one of the three modules does not work, modified marks cannot be spread orderly, leading to the epigenetic instability that would further lead to pathological problems. For example, if failing to propagate to offspring due to abnormal gene expression pattern or defective replication or mutations in modification enzymes, the epigenetic information would lead to irregular developmental programs and event to tumorigenesis, cancer, cellular senescence and apoptosis [65]. Our multiscale model can well reveal the essential mechanism of the synergism even in a more realistic case. In particular, our result on the synergism indicates that through the synergism among histone modification, chromatin organization and gene transcription, can we manage and explain complex mechanisms of genetic and epigenetic regulations. This result may shed light on functional mechanisms, which provide useful clues for experiments in the future.

We emphasize that our multiscale model is also a useful approximation in study of chromatin dynamics. Specifically, we modeled chromatin as a polymer and used a generalized Rouse model to describe the polymer dynamics. Recall that Rouse-type models such as SBS model [66], Rod-like model, Zimm model, reptation model [67] can also represent a self-avoiding polymer. However, the Rouse model is suitable for the situation where the environmental effects of entanglement and crowding are negligible [68]. When modeling the processes of nucleosome modifications and gene transcription, we used a coupled reaction system suitable to the use of the Gillespie algorithm [51]. This implies that we have made the Markovian assumption, that is, the stochastic motion of enzymes is uninfluenced by previous states, only by the current state. But, in vivo, intracellular biochemical processes occur, in general, in a memory manner, leading to non-Markovian kinetics [69]. In spite of the Markov assumption, our model can also be extended to non-Markovian cases.

Recently, the work of Michieletto *et al.* [27] on epigenetic recoloring dynamics based on the potential of the whole system revealed a pathway for the epigenetic information establishment and heritability. However, their approach cannot accurately dissect the contribution of 1D and 3D coupling to epigenetic dynamics, thus failing to stress the effects of long-range interaction caused by the chromatin dynamics. The work of Jost *et al.* [28] based on a LC model stressed the importance of long-range interaction or chromatin conformation in epigenetic maintenance, but the proposed method did not consider time explicitly, failing to describe the dynamic processes of chromatin configuration and epigenetic changes in multiple time scales. In contrast, our model explicitly considered gene transcription and DNA replication, and provided an effective framework for analyzing the relationships among epigenetic maintenance, chromatin configuration and gene transcription.

In summary, our model provides a study paradigm for 4D nuclear project even in a more realistic or complex case. Our findings, which rationalize the mutual effects of spatial folding, epigenetic modification and gene function on the establishment and maintenance of stable epigenetic cell memory, provide useful clues for experiments on the impacts of conditions related to epigenetic chromatin such as histone exchange [16,70–72], cancer therapy [73], apoptosis [74].

## Notes

### Competing Interest Statement

The authors have declared no competing interest.

