## Supplemental File for "Synergism between Chromatin Dynamics and Gene Transcription Enhances Robustness and Stability of Epigenetic Cell Memory"

**This PDF file includes:**

Supplementary Notes 1 to 3.

Supplementary Table 1.

Supplementary Figures 1 to 4.

Supplementary Movies 1 to 4

References

**Other Supplementary Material for this manuscript includes the following:**

(available at XXX)

Movies S1 to S4

Simulation codes

### Supplementary Note 1: Reasonable simplification of genetic and epigenetic processes

Pervious works of modelling genetic and epigenetic processes focused, primarily, on how chromatin domains are organized, and did not simultaneously consider chromatin structure, epigenetic modifications and gene transcription (or gene activity) nor the relationship among them. Based on biological reality, here we build a multiscale model for this relationship. Our model considers three modules as specified in the main text. Specifically, for chromatin structure, we model chromatin as a chain consists of finite monomers (or beads) with each representing a nucleosome with a 3-dimensional (3D) position vector. For epigenetic regulation, we only consider three kinds of nucleosome states - methylated (M), unmodified (U), and acetylated (A). However, it should be pointed out that, in vivo, the process of epigenetic modification would be much more complex in contrast to the situation considered here. For gene transcription, we only consider the manner of constitutive expression and do not consider that of bursty expression – a popular gene expression manner in eukaryotic cells. For these simplifications, we will give some interpretations in the following.

***Modification process***

Nucleosome is the basic unit of chromatin with DNA wound around histone octamer surface like thread around a spool. Each nucleosome has two copies of a histone, and biochemical modifications occur on histone amino (N)-terminal tails. However, our model ignores the role of "hetero-modified" nucleosomes [1] and only considers modifications at the nucleosome level. In vivo and in vitro, there exist mono-, di-, and trimethylated forms such as H3K27me1, H3K27me2 and H3K27me3. In most situations, it is not clear whether multiple methylation reactions occur at a time or in a separate step, but it is well known that a methylation reaction can occur in a processive or distributive manner [2]. For the sake of simplicity, we ignore the case of monomethylation or dimethylation, and only consider that a protein complex can effectively add or remove three methyl groups in one step.

We point out that not all methylations suppress gene expression. For example, H3K4me1 in enhancers activates transcription, whereas H3K36me and H3K79me in the transcribed region affect transcriptional elongation [3]. Some nucleosomes such as the H3K9me, H3K27me, H4K20 and other post-transcriptionally methylated nucleosomes play an important role not only in repressing gene expression but also in regulating and mediating the timing of cellular activities [3]. Our model makes the assumption – the methylated nucleosomes are linked to a condensed structure that hinder transcription – because methylation and acetylation on the same nucleosome such as H3K9 and H3K27 have distinct states. This assumption is for convenience of comparison with acetylation.

***Transcription***

In prokaryotes and eukaryotes, transcription is a complex process and takes place often in a bursty manner. In our multiscale model (see the main text), due to the complexity and uncertainty of transcription machinery as well as our interest manly in the relationship between transcription (in fact, gene activity) and chromatin dynamics, we use a reaction to model a transcription process without considering details of transcriptional burst. Therefore, we posit that the occurrence of transcription events depends on the current chromatin state and that transcription can happen anywhere within the gene.

***DNA replication and mitosis***

The mechanism of nucleosome modification inheritance through DNA replication remains unclear. In any working model on nucleosome modification inheritance, the epigenetic information needs to be copied from parental nucleosomes to newly synthesized ones [4]. In our model, we suppose that de novo nucleosomes must be participated in the two copies of DNA, and both the old and new nucleosomes are normally yet randomly shared by two daughter chromosomes.

DNA is condensed from molecules to chromosomes during mitotic prophase. It ceases to function as accessible genetic material (transcription stops) and becomes a compact transportable form [5]. After DNA replication and at the beginning of next cell cycle, chromosomes unwind into chromatin. Therefore, we can assume that the effect of chromosome during mitosis is so low or even none that mitosis may be neglected or may not be considered.

### Supplementary Note 2: Mathematical modeling of modification-mediated chromatin dynamics

Our multiscale model with fast and slow timescales, which describes the epigenetic dynamics of chromatin as specified in the main text, involves the decoding of epigenetic mechanisms consisting of chromatin modifications and configurations as well as gene activities, which are related to memory and switching. In our model, the 3D organization of chromatin might significantly impact epigenetic regulation via local and long-range interactions between nucleosomes (seeing the main text). The 3D shape evolves by the total potential of the system mediated by modification states. Methylated nucleosomes are self-attractive, which would compact the chromatin and lessen the binding of transcriptional factors. And acetylated state is self-repulsive, which would enhance global looseness and accelerate the transcription that in turn would facilitate the demethylase and acetylation processes. Nucleosome modification operates on the 3D space through updating the energy potential of the system to regulate the chromatin motion. Next, we will make detailed supplementary explanations for our mathematical model.

***Generalized Rouse model***

We use a generalized Rouse model with additional interacting force to describe the chromatin organization. Rouse chain is consisting of a collection of beads connected by harmonic springs (spring constant is $\kappa$). The Rouse model neglects the contributions of volume exclusion and hydrodynamic interactions, and assumes the diffusion coefficient $D$ is independent of the position of each monomer. In addition, because of the fact that Rouse model alone cannot completely reflect the different states of chromatin in different modifications, we add some attraction / repulsion forces according to previous results [6]. M state is self-attractive and we let $\kappa_M = \kappa/1000$ represent the attractive interaction coefficient, A state is self-repulsive and we let $\kappa_M = -\kappa/30000$ represent the repulsive interaction coefficient, while the U state is a neutral state that does not self-attract but can participate in epigenetic dynamics. Note that the interaction coefficient between the modified nucleosomes ($\kappa_M, \kappa_A$) is much smaller than the spring stiffness between adjacent monomers ($\kappa$). Therefore, we use the generalized Rouse Model to describe the conformational dynamics of the polymer and use the Brownian motion to represent the chain diffusion.

Using the above interaction coefficient, we test the radius of gyrations of different modification states with 60 nucleosomes (Supplementary Fig. 1a). When all the 60 nucleosomes are acetylated

(A), the radius of gyration is about $8.5\mu m$. When all the 60 nucleosomes are methylated (M), the radius of gyration is about $2.5\mu m$. When all the 60 nucleosomes are unmodified (U), the radius of gyration is about $5.5\mu m$. Distinct levels of modification show different 3D chromatin structures, which enable us to better analyze the relationship between structure and function.

Based on the above analysis, for the epigenetic propagation, it is reasonable to set different interaction gyration with different modification: acetylated (methylated) monomer have the larger (smaller) spatial interaction gyration. We consider the radius of space interaction influence of a monomer: when the distance is smaller than a certain threshold $d_S$ with $S \in \{M, A\}$, the monomer is able to spread its mark; when the distance is larger than this threshold, the monomer is also able to spread, but with a reduced efficacy like a power function with decay exponent $-3/2$ (Supplementary Fig. 1b). We set the interaction distance threshold as $d_M = b/2.4$ or $d_A = b/1.2$, where $b$ is the standard deviation (STD) of distance between adjacent monomers.

In order to test the rationality of the above hypothesis, we calculated the average numbers of methylated nucleosomes and acetylated nucleosomes around an unmodified nucleosome respectively (Supplementary Fig. 1c). Then, we find that for stable memory, the average numbers of the nucleosomes participated in modifications are approximately equal ($\approx 8.5$) in the distinct modification states.

***Biochemical reaction system & Gillespie algorithm***

In our system of $N$ nucleosomes, there are in total $4N+1$ possible pathways, each consisting of $N$ nucleosome methylations, $N$ nucleosome demethylations, $N$ nucleosome acetylations, $N$ nucleosome deacetylations, and transcription. Stochastic simulations of those reactions are carried out using the Gillespie algorithm. The algorithm is defined by a series of biochemical reactions and a relevant propensity for each biochemical pathway to occur. At each iteration, it generates probabilistically a time-step $\tau$ representing when the next reaction occurs and a reaction-index $a$ representing which reaction will occur. In the time-step $\tau$, the chromatin updates the position by the Brownian motion. When the system time increases by $\tau$, the selected reaction $a$ is then performed by updating the system state.

This algorithm can generate not only exact sample paths based on probability distribution but also roughly accurate time step corresponding to the typical time-scale of modification and transcription. In order to test this, we calculate the probability distribution of time steps in three bins - $(0,60]$ (second), $(60,3600]$ (minute) and $(3600, \infty)$ (hour) (Supplementary Fig. 1d). Then, we find that the probabilities in three bins are 0.18, 0.78 and 0.04. Most time steps fall to the minute timescale. Thus, it is reasonable and appropriate to use the Gillespie algorithm to simulate a genetic and epigenetic regulation networks.

### Supplementary Note 3: Simulations and quantitative analysis

For a given set of parameter values, the equilibration of 5 cell cycles was run before taking samples of the system and this was to ensure conformational and epigenetic equilibration. After that, snapshots of the system were taken every 300 seconds. In addition to the quantities mentioned in the main text (epigenetic magnetization, radius of gyration and gene activity), we also calculated some other quantities.

***Bistability measurement***

The quantity introduced in [7] used to determine the probability for the system to be in one of three epigenetic states is $P_M$, the probability that the number of methylated nucleosomes exceeds that of acetylated marks ($s > 0$), expressed by

$$P_M = \Pr(s > 0).$$

Similarly, we have another probability expressed by

$$P_A = \Pr(s < 0).$$

Thus, bistability can be measured according to

$$B = 4P_M P_A$$

Note that if $P_M = P_A = 0.5$, quantity $B$ is equal to 1. Therefore, when $B$ is close to 1, this means the occurrence of bistability.

***Mean first passage time***

First passage times (FPT) $t_{FP(M)}$ and $t_{FP(A)}$ are defined as the times for the system to switch to the opposite state, when initialized in the uniform M or A state, respectively. For example, for a fully repressed state $s = 1$, the FPT is the one that the epigenetic magnetization changes to $s < 0$. That is

$$t_{FP(M)} = \min(t|s < 0)$$

Similarly,

$$t_{FP(A)} = \min(t|s > 0)$$

We point out that in our demonstration of numerical results, the MFPT is an average value obtained after many simulations.

Since a nucleosome that is randomly inserted during DNA replication is an unmodified state, it is necessary to allow the system to recover from a perturbation before assessing the stability of the state after DNA replication. For this reason, results of bistability measurements and MFPT are obtained only by calculating the last snapshot of each cell cycle.

### Supplementary Table 1: List of model parameter values

| Parameter | Description | Value |
|---|---|---|
| | Chromatin motion | |
| $d$ | dimension | 3 |
| $\kappa$ | spring constant $N/\mu m$ | $dD/b^2$ [8] |
| $\kappa_M$ | attraction interaction constant $N/\mu m$ | $k/1000$ |
| $\kappa_A$ | repulsion interaction constant $N/\mu m$ | $-k/35000$ |
| $d_M$ | interaction distance $\mu m$ | $b/2.4$ |
| $d_A$ | interaction distance $\mu m$ | $b/1.2$ |
| $b$ | STD of distance between adjacent monomers $\mu m$ | $\sqrt{3}$ |
| $D$ | monomer diffusion coefficient $\mu m^2/s$ | 1 |
| $\Delta t$ | time step $s$ | 0.01 [6] |
| | Chromatin modification | |
| $k_{me}$ | methylation rate (U to M) $nucleosome^{-1}s^{-1}$ | free |
| $k_{ac}$ | acetylation rate (U to A) $nucleosome^{-1}s^{-1}$ | free |
| $k_{dem}$ | demethylation rate (M to A) $nucleosome^{-1}s^{-1}$. | $k_{ac}/10$ |
| $k_{dea}$ | deacetylation rate (A to M) $nucleosome^{-1}s^{-1}$ | $k_{me}/10$ |
| $\gamma_{me}$ | noisy methylation rate (U to M) $nucleosome^{-1}s^{-1}$ | $k_{me}/20$ |
| $\gamma_{ac}$ | noisy acetylation rate (U to A) $nucleosome^{-1}s^{-1}$ | $k_{ac}/20$ |
| $\gamma_{dem}$ | noisy demethylation rate (M to A) $nucleosome^{-1}s^{-1}$ | $k_{me}/20$ |
| $\gamma_{dea}$ | noisy deacetylation rate (A to M) $nucleosome^{-1}s^{-1}$ | $k_{dea}/20$ |
| | Transcription | |
| $p_{dem/ac}$ | demethylation/acetylation probability $nucleosome^{-1}transcription^{-1}$ | 4e-3[9] |
| $\mu_{\min}$ | minimum transcription initiation rate $transcription \cdot s^{-1}$ | 1e-4[9] |
| $\mu_{\max}$ | maximum transcription initiation rate $transcription \cdot s^{-1}$ | 4e-3 |
| | Other parameters | |
| $N$ | number of nucleosome | 60 |
| | cell cycle length (h) | 22 [10] |

### Supplementary Figure 1

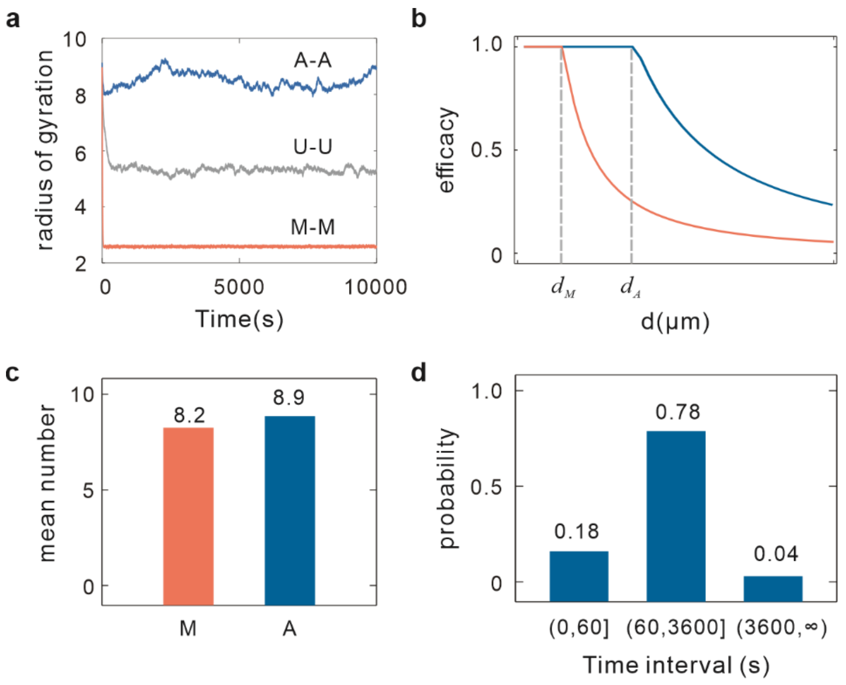


**Supplementary Fig. 1** (a) Radius of gyration in different states. (b) The function of impact factor $IF_j^S, j \in X_i, S \in \{M, A\}$ with respect to interaction spatial distance. (c) The average numbers of methylated nucleosomes and acetylated nucleosomes around an unmodified nucleosome. (d) Histogram of time steps generated by the Gillespie algorithm.

### Supplementary Figure 2

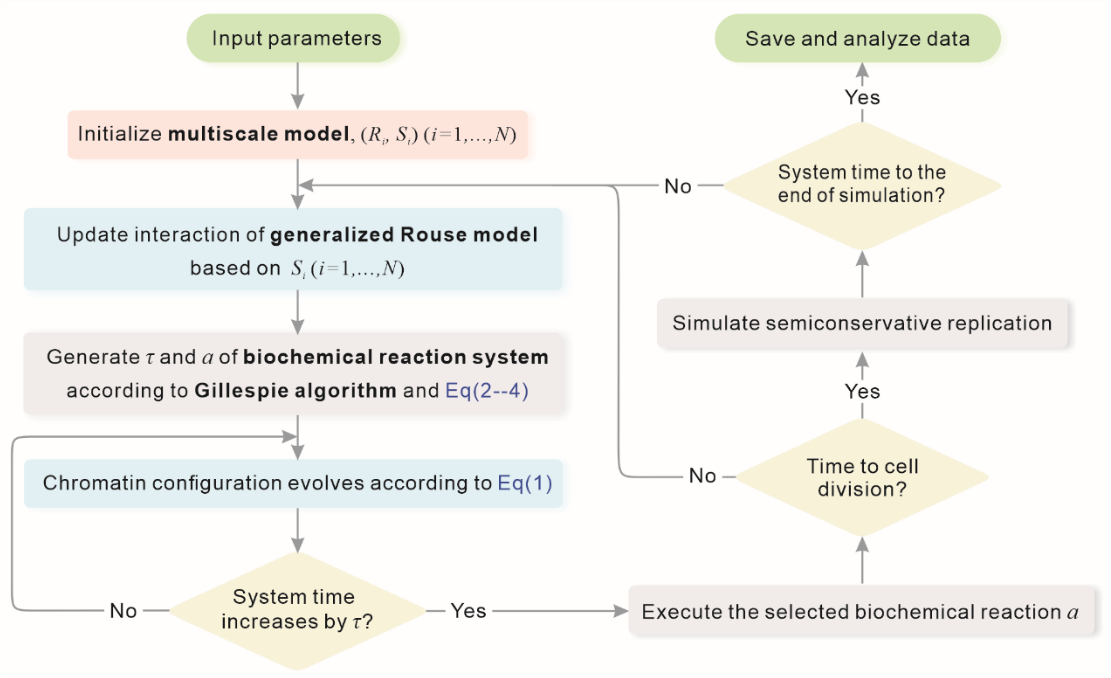


**Supplementary Fig. 2 Flowchart for simulation of the multiscale model.**

### Supplementary Figure 3

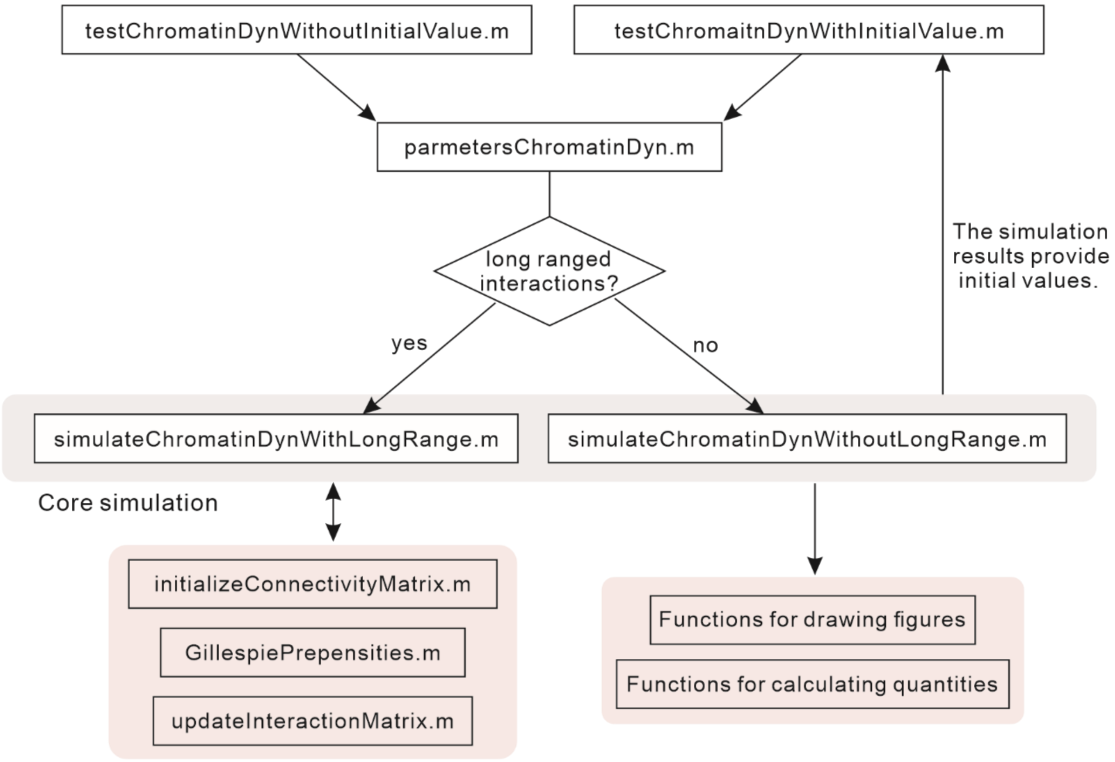


**Supplementary Fig. 3 Flowchart for our program codes.**

### Supplementary Figure 4

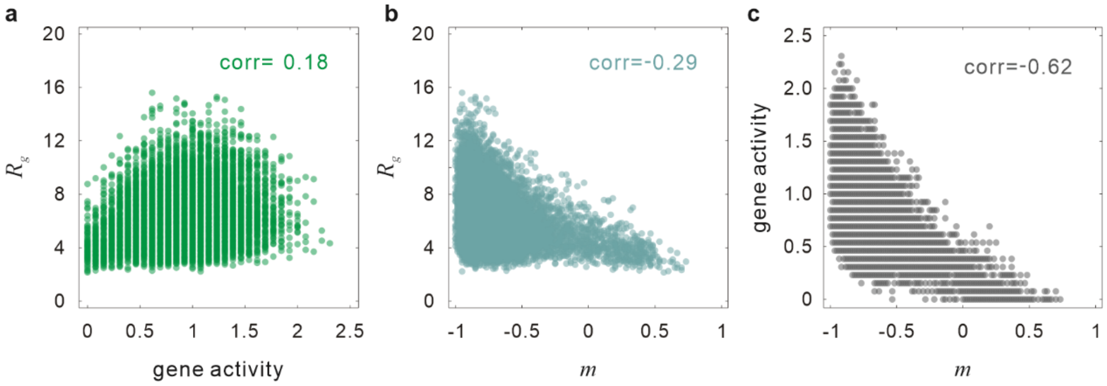


**Supplementary Fig. 4 Shown are the Pearson correlation coefficients between the radius of gyration $R_g$ and gene activity (a), between $R_g$ and magnetization $m$ (b), and between gene activity and $m$ (c).** Comparing this figure and Fig. 5 in the main text, we can clearly see that the long-range interaction can better maintain the stability of two alternative states than the local interaction, and that the correlations in the former case are higher than those in the latter case.

### Supplementary Movies 1 to 4

**Movie 1**: A simple case for polymer motion with random chromatin modification state and position. The sampling timescale of chromatin folding is $10^{-2}$ s and the total time is 1 s. In this second, the modified state of chromatin does not change.

**Movie 2**: Simulation of a polymer with stable methylated states. Snapshots of the system are taken every 300 seconds and data for one cell cycle are recorded to produce the movie.

**Movie 3**: Simulation of a polymer with stable acetylated states. Snapshots of the system are taken every 300 seconds and data for one cell cycle are recorded to produce the movie.

**Movie 4**: Simulation of the process from stable methylation state to stable acetylation state. Snapshots of the system are taken every 300 seconds and data for ten cell cycles are recorded to produce the movie.
